# Dynamic co-expression modular network analysis of bHLH transcription factors regulation in the potato spindle tuber viroid-tomato pathosystem

**DOI:** 10.1101/2023.11.10.566618

**Authors:** Katia Aviña-Padilla, Octavio Zambada-Moreno, Marco A. Jimenez-Limas, Rosemarie W. Hammond, Maribel Hernández-Rosales

**Affiliations:** Centro de Investigación y de Estudios Avanzados del I.P.N Unidad Irapuato; Department of Crop Sciences, University of Illinois at Urbana–Champaign, Urbana, Illinois; Centro de Investigación en Computación, Instituto Politécnico Nacional, Ciudad de México; USDA, Agricultural Research Service, Beltsville Agricultural Research Center, Beltsville Maryland

**Keywords:** viroids, plant-pathogens, co-expression, gene modules, biological networks, bHLH-TF, computational biology, systems biology

## Abstract

Viroids, minimalist plant pathogens, present significant threats to crops by causing severe diseases. The use of high-throughput sequencing technologies for analyzing the transcriptomes of viroid-infected host plants has yielded informative information on gene regulation by these pathogens, however a complete understanding of the transcriptome data suffers from the inclusion of numerous genes of unknown function. Co-expression analysis addresses this by clustering genes into modules based on global gene expression levels. Our previous study emphasized basic helix-loop-helix protein (bHLH) transcriptional reprogramming in tomato in response to different potato spindle tuber viroid (PSTVd) strains. In the current research, we delve into tissue-specific gene modules, particularly in root and leaf tissues, governed by bHLH transcription factors during PSTVd infections. Utilizing public datasets that span Control (C; (mock-inoculated), PSTVd-mild (M), and PSTVd-severe (S23) strains in time-course infections, we uncovered differentially expressed gene modules. These modules were functionally characterized, identifying essential hub genes. We identified the roles of bHLH transcription factors (TFs) in managing processes like photosynthesis and rapid membrane repair in infected roots. In leaves, external layer alterations influenced photosynthesis, linking bHLH TFs to distinct metabolic functions. Expanding on these findings, we explored bipartite networks, discerning both common and unique bHLH TF regulatory roles, notably highlighting the bifan motif’s significance in these interactions. Through this holistic approach, we deepen our understanding of viroid-host interactions and the intricate regulatory mechanisms underpinning them.

## Introduction

Viroids are small, infectious RNAs that cause economically important diseases in plants [1,2]. Remarkably, despite their tiny size of 246–401 nucleotides, they have a distinctive single-stranded circular RNA genome that operates devoid of coding capacity [3]. Operating independently, these minuscule agents replicate within plants by leveraging the host’s enzymatic machinery [4]. Beyond their impact on crops, viroids serve as a valuable model for unravelling intricate host-pathogen interactions, extending the scope of exploration into biological dynamics [5–9]. This multifaceted nature positions viroids not only as plant pathogens but allows further exploration into the depths of biological processes [10]. In essence, viroids offer insights into molecular and systems biology host-pathogen interplay, providing a distinctive perspective [11].

Viroid infections have significant repercussions as they disrupt host development and interfere with crucial physiological processes [11–18]. An illustrative instance can be observed in tomato, a vital global agricultural commodity that yielded more than 189 million tons in 2021 (http://faostat.fao.org/, accessed in July 2023). This staple crop faces a tangible threat from viroids, underscoring the potential impact of these infectious agents on essential food production systems. Emerging reports link viroids to hormone pathways and transcription factor dynamics, disrupting plant gene expression landscapes [19]. Transcriptional profiling has been instrumental in understanding the global effects of viroid infections on plant gene expression.

High-throughput technologies have enabled comprehensive studies of viroid-plant interactions, shedding light on gene expression patterns on a large scale [10]. Yet, the presence of unknown genes in plant genomes poses challenges in data interpretation. Co-expression network analysis addresses this issue by clustering genes based on their expression levels [20–21]. This network forms a graph of interconnected nodes representing genes with similar expression, enabling insights into molecular pathways [22]. Constructing co-expression networks across diverse conditions, such as disease states, facilitates the identification of disease-induced changes through network connectivity patterns [23–29]. Analyzing modular gene co-expression within these networks unveils the systemic functionality of genes [23, 24–29].

Topological features within networks reveal key molecular components in complex biological systems, including gene co-expression relationships [30]. Modular gene co-expression analysis is a potent method for uncovering the broader functionalities of genes at a systems level [23]. By deciphering network modules, insights into gene interactions and potential roles can be gleaned [21–25]. Notably, genes with shared functions often exhibit robust correlations in their expression levels, laying the foundation for uncovering molecular pathways underlying diseases and conditions [31–33].

Genetic and molecular studies centered on plant transcription factors (TFs) have provided invaluable insights into plant-specific responses, encompassing defense against pathogens, responses to light, and environmental stresses like cold, drought, high salinity, and developmental processes [34–36]. Transcriptional regulation, a fundamental process governing tissue-specific gene expression and responses to stimuli, underlies these processes [37]. Among the key regulators of gene expression, TFs bind DNA sequences to fine-tune transcription, with distinct strategies across different life forms [35–37].

In a previous study, we introduced a comprehensive tomato gene co-expression network interwoven with hormone pathways, shedding light on gene expression during potato spindle tuber viroid (PSTVd) variant infections [38]. The network showcased the involvement of Basic-Helix-Loop-Helix (bHLH) biomolecules. bHLH transcription factors (TFs) were engaged in regulatory interactions, notably distinct in mild- and severe-viroid-infected samples, hinting at their potential significance in the progression of viroid diseases. bHLH members were highlighted as master regulators in the PSTVd-tomato pathosystem, underlining their significance [38]. Hence, we selected bHLH TFs to further investigate guided regulatory transcriptional events.

bHLH TFs, prevalent in eukaryotes, constitute a substantial TF family. Anchored by a DNA-binding bHLH domain, these proteins orchestrate various biological processes, including stress responses [36,39]. The bHLH domain’s N-terminus contains a DNA-binding motif, with proteins binding to sequences harboring the E-box (59-CANNTG-39) consensus motif, commonly the G-box (59-CACGTG-39). Plant bHLH TFs play diverse roles, including hormone signaling, stress responses, and cellular differentiation.

This study aims to unveil co-expression modules of the bHLH family, highlighting their roles in the tomato immune response against viroid infection. Focusing on bHLH TFs as hub genes, we explore their involvement in host molecular mechanisms. Our main goal is to identify conserved and distinct bHLH-guided transcriptional programs using systems biology approaches.

Our results reveal that bHLH TFs regulate photosynthesis and membrane lipid repair in infected roots. Furthermore, our findings shed light on the intricate regulatory networks orchestrated by bHLH TFs in leaf tissue during PSTVd variant infections. The disruptions in cuticle function, shifts in metabolic processes, and alterations in biosynthesis pathways portray the plant’s adaptive response to mitigate viroid impact. Notably, our findings emphasize the enrichment of the bifan motif, a critical regulatory interaction, conserved and intricately woven across the analyzed biological networks. This comprehensive understanding contributes to unravelling the host-pathogen interaction and may provide strategies to enhance plant resilience against viroid infections.

## Materials and Methods

### 2.1. Designed Pipeline for the Networks Approaches

We performed a comprehensive comparison of the effects exerted by PSTVd (M; mild) and (S23; severe) strains on the susceptibility of the tomato host, with a primary focus on symptom development. To achieve this, we employed an innovative approach that integrates transcriptomic and functional genomics data. Through this strategy we implemented co-expression and module network analysis, as depicted in **Figure 1**. The aim was to unveil the intricate panorama of bHLH-guided transcriptional events governing the *Solanum lycopersicum* response to PSTVd infection. Our designed codes are available at https://github.com/kap8416/Dynamic-coexpression-modular-analysis-BHLH-in-PSTVD-tomato. Our methodology can be delineated into a series of crucial steps. First, we conducted a tissue-specific module network analysis with a special emphasis on the bHLH interactions (Steps a–b). Subsequently, the CEMiTool package, [40] performed the inference of gene co-expression modules (Step c). Then, employing the output files derived from the CEMiTool analysis, we proceeded to construct bipartite networks for each tissue (Step d). Finally, we explored representative motifs within the modules (Step e), unveiling essential regulatory motifs that underscore the underlying dynamics.

**Figure 1.**
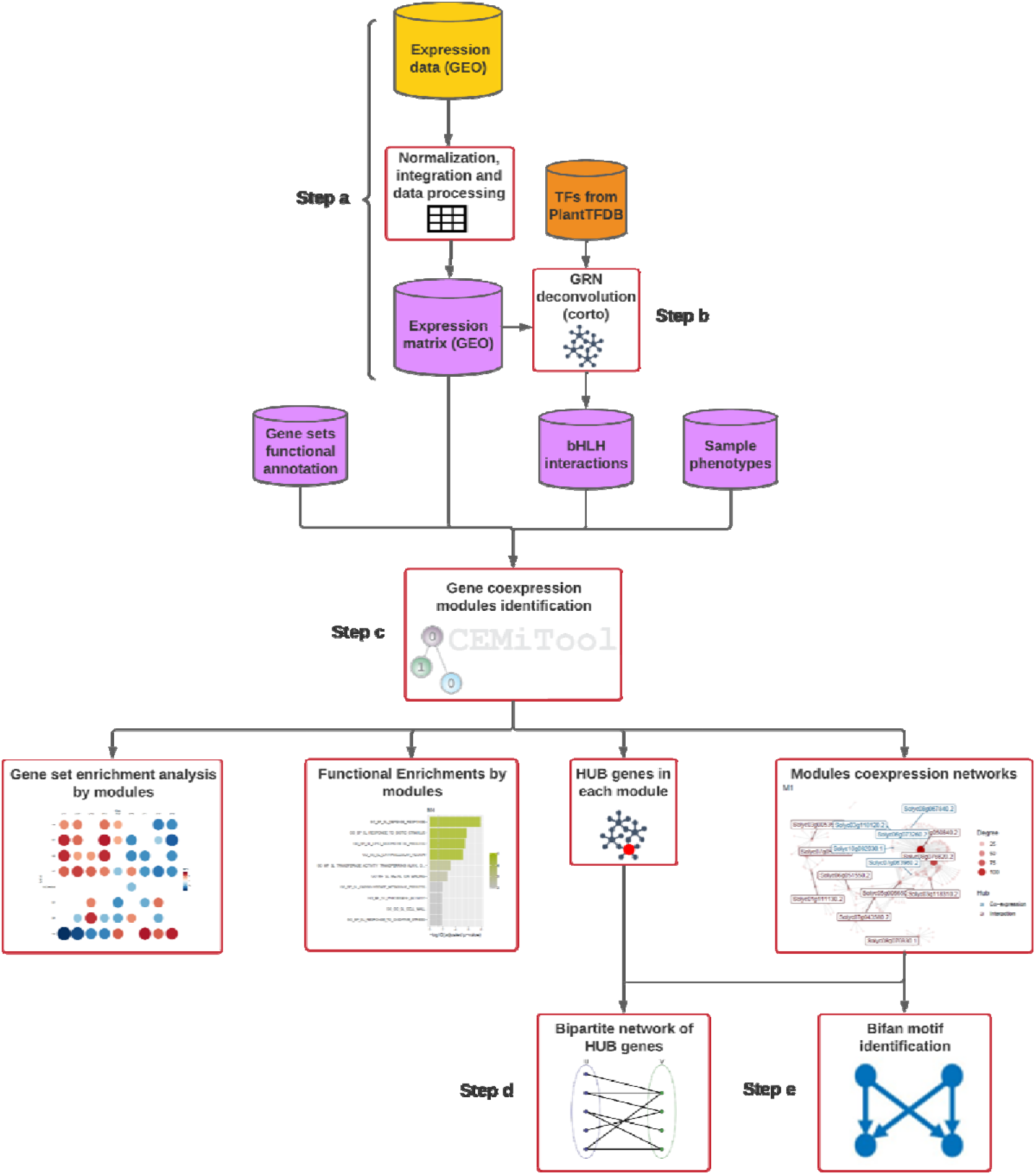
Pipeline for the networks approach. (a) Commencing the process, we acquired publicly available microarray expression data encompassing both control and infected leaves and roots.(b) With the obtained expression data, we constructed an expression matrix of each tissue (roots, and leaf separately) and curated additional pertinent files, collectively serving as the foundational input for our subsequent analysis: functional annotation, expression matrix, interest genes interactions, and sample phenotypes (the required input files are depicted in pink). (c) We performed a co-expression modular network analysis. This pivotal step was facilitated by the utilization of the CEMiTool, a specialized tool for inferring co-expression modules. (d) Following the completion of the network analysis, we delved into a series of pivotal explorations. Gene Set Enrichment Analysis (GSEA) was performed to unveil significant associations, and functional enrichment analysis was executed on the modules. Furthermore, we meticulously identified the hub genes residing within the network clusters. (e) With the elucidation of crucial insights from the previous steps, our focus shifted towards the creation of bipartite networks delineating the bHLH-regulon interactions. (g) Using our comprehensive network structures, we conducted a final phase of exploration. We identified and investigated the most representative motifs within each network module.

### 2.2. Description of the Gene expression Datasets

Two expression matrices depicting the comprehensive effects of PSTVd infection on the tomato plants’ transcriptome were utilized. Microarray datasets from the roots (GSE111736) and leaves (GSE106912) of tomato plants under control (C), mild infection (M), and severe infection (S) conditions were obtained from the NCBI Gene Expression Omnibus database (https://www.ncbi.nlm.nih.gov/gds, accessed on 4 June 2022). These studies employed a time-course analysis spanning three distinct stages: early symptoms (17 dpi), pronounced symptoms (24 dpi), and recovery (49 dpi), resulting in a total of 26 root and 27 leaf samples being individually assessed [41, 42].

### 2.2. Obtaining the bHLH-gene specific interactions

To construct gene co-expression networks corresponding to each module, we utilized a compilation of interactions involving the bHLH transcription factors (TFs). This compilation was derived from a prior study conducted by our team, wherein gene regulation networks (GRNs) specific to the tomato-PSTVd interaction were established. Spearman correlation coefficients were computed using the Corto R package to facilitate this process. Subsequently, a subnetwork was extracted from this GRN, focusing exclusively on the bHLH TFs and their associated interactors. The detailed methodology for constructing this network is available in Aviña-Padilla et al., 2022, [38].

### 2.3. Dynamic co-expression modular network analysis

For the identification of gene co-expression modules, we utilized the CEMiTool webtool [citation to CEMITOOL]. This comprehensive package employs a systems biology approach, offering seamless and automated analysis of modules. The modular analysis was conducted by inputting the following datasets (**Figure 1c**):

- A tab-delimited text gene expression matrix customized for each tissue.
- A tab-delimited text table containing interaction details of the bHLH-TFs.
- Tomato gene ontologies presented in `GMT` file format.
- A text file table with “SampleName” and “Class” columns, providing descriptive information about the phenotype of each sample.

The analysis was executed with default parameter settings. Prior to analysis, the expression matrices underwent normalization using the RMA function from the `limma R` package, [43]. This normalization aimed to derive significant Beta-values crucial for accurately inferring co-expression modules.

### 2.4. Obtaining bHLH-regulon bipartite networks and functional assignment

Based on the outcomes of the modular co-expression analysis, we constructed bipartite networks for the most intriguing co-expression modules pertaining to each tissue. These networks were created utilizing `Python` scripts. Within these visualizations, all the bHLH-TFs involved in each tissue were represented, alongside the genes they regulate (referred to as the regulon). These components were categorized based on the way their regulation occurs. Moreover, we pursued functional enrichment analysis by employing gene ontology terms for distinct categories of regulation. To execute this step, we employed the `Gprofiler` package [44] within the RStudio environment. This enabled us to discern and characterize the functional attributes associated with the regulatory dynamics observed within the bipartite networks.

### 2.5. Identifying Regulatory Motifs within Network Modules

After obtaining the network modules for both leaf and root tissues, we searched for significant regulatory motifs within each of these modules. To achieve this, we utilized the `Mfinder` tool, a specialized resource designed for detecting network motifs. This tool employs two distinct methods for motif identification: 1) Full Enumeration of Subgraphs: This technique involves an exhaustive enumeration of subgraphs within the network. The concept of network motifs, seen as fundamental building blocks of complex networks, forms the basis of this approach [45]; 2) Sampling of Subgraphs for Concentration Estimation: in this method, subgraphs are sampled to estimate their concentrations within the network. By assessing prevalence and significance, this approach aids in detecting network motifs [46]. To execute this process, a `.txt file` containing the module of interest was submitted to the `Mfinder` tool. The tool’s output comprises significant motifs identified within the network. This step enables the discovery of regulatory interaction patterns that play a pivotal role in shaping the gene co-expression networks.

## Results

### 3.1 Gene Co-expression Modules and Their Distinct Regulatory Patterns in Root Tissue

In our exploration of root tissue, we identified 10 distinct gene co-expression modules, designated as M1 to M10, with varying sizes ranging from 216 to 23 genes. We selected the modules that showcased distinctive regulatory dynamics and potential functional implications across the different experimental conditions, as depicted in **Figure 2**.

**Figure 2.**
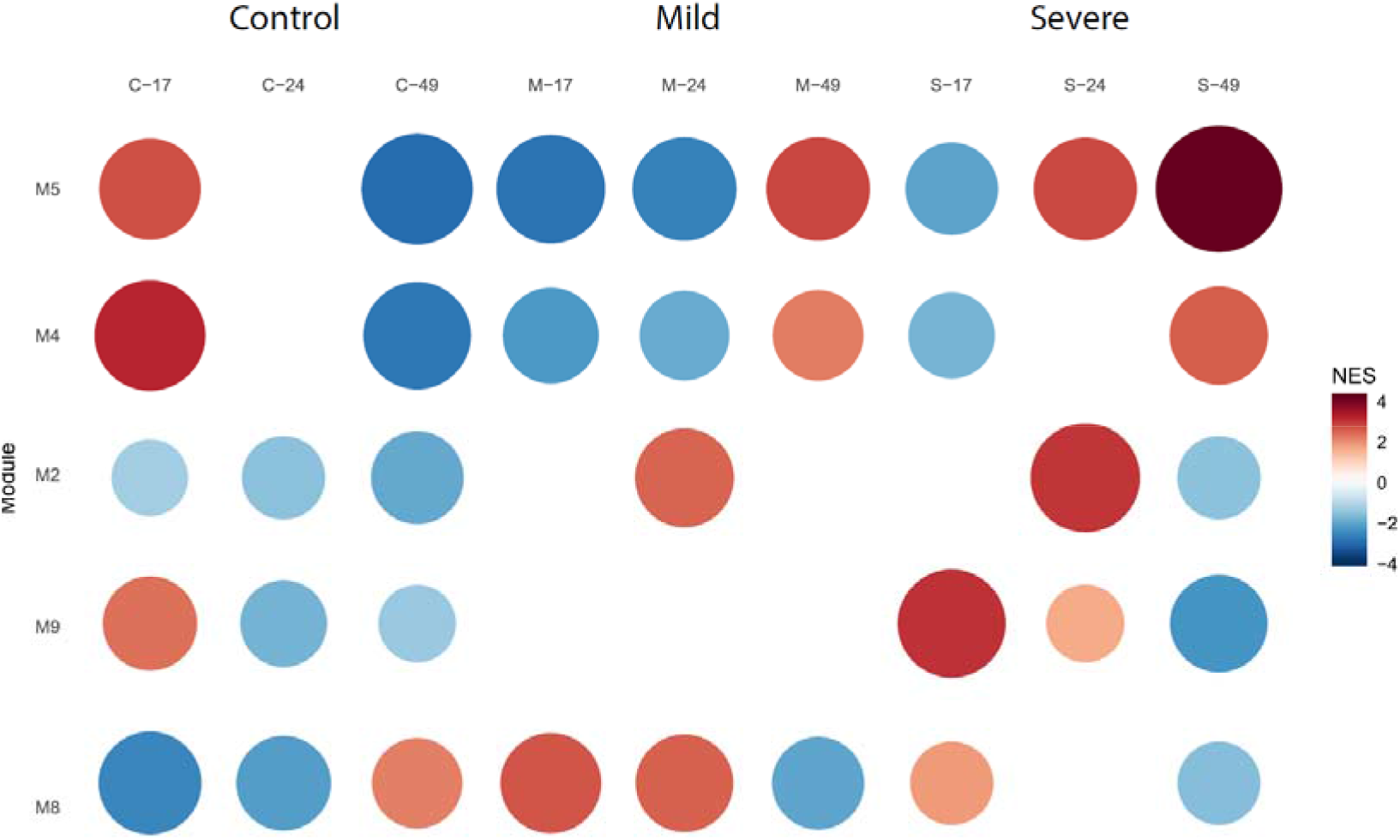
Dynamics of Co-Expression Modules in Root Tissue. Gene Set Enrichment Analysis (GSEA) was employed to unveil the module activity across distinct conditions, including control, PSTVd M, and PSTVd S23 strains, throughout the time course analysis of root tissue. Notably, the M5, M4, M2, M9, and M8 modules emerged as highly significant in this context, showcasing distinctive regulatory dynamics and potential functional implications across the different experimental conditions.

#### 3.1.1 Modules that Exhibit Strong Symptom Induction in Response to the Severe Strain

Three modules, M2, M4, and M5, displayed notable regulatory changes during the 24-day symptom development stage of the severe strain, as depicted in **Figure 3**. These modules show a synchronized spike in gene expression as the symptoms intensify, indicating the plant’s heightened response to the severe PSTVd strain. This surge in gene activity highlights the plant’s determined defense efforts against the aggressive pathogen. Such observations provide insights into the plant’s adaptive defense tactics against the severe strain, aiding its survival and recovery.

**Figure 3.**
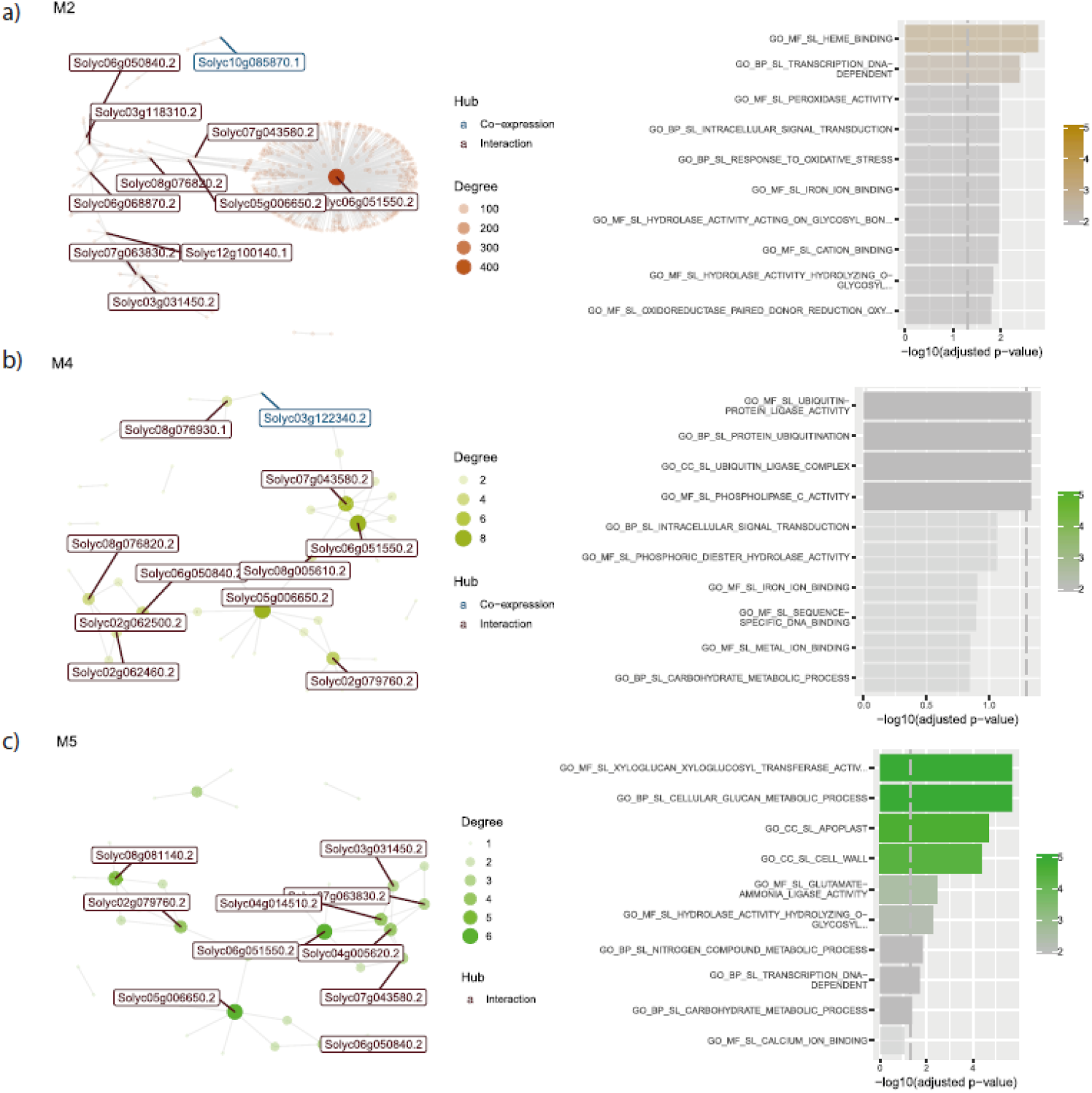
Co-expression interaction network and functional enrichment of modules firstly repressed and then overexpressed in root. Each node on the network represents a gene, size is proportional to the amount of interactions (degree). Hub genes are listed with their names, in red if they only interact, and in blue if they are co-expressed with their targets. Next to each network, biological functions of their genes are shown. Length of the bar represents significance, and color gradient number of genes implicated in each Gene Ontology Term. a) Interaction network and biological functions for M2; b) M4; and c) M5.

##### 3.1.1.1 Module 2: Coordinated Responses associated to (fe inefficient) bHLH transcriptional regulator

Module 2, comprising 123 genes, underscores the integral role of the bHLH transcriptional regulator, *Solyc06g051550*, in modulating iron uptake efficiency, this TF present interaction with more than 400 hundred genes in this module, which points out its importance. The Gene Set Enrichment Analysis (GSEA) as illustrated in **Figure 2**, denotes a marked upregulation during symptom proliferation in both PSTVd infection intensities, implying Module 2’s cardinal role throughout the infection trajectory. Notably, this regulation is predominantly discernible within the epidermal cells and marginally within the outer cortical cell strata of root apexes. The gene regulator exhibits an affinity for heme binding, hinting at its potential involvement in iron metabolism regulation. Simultaneously, *Solyc10g085870.1* (depicted in blue in **Figure 3a**) is co-expressed in the network. This gene encodes a UDP-glycosyltransferase 73C3 that plays a crucial role in glycosylation processes, pivotal for modulating the metabolism of myriad cellular compounds.

##### 3.1.1.2 Module 4: Lipoxygenase D as a Key Player in Defense Processes

Module 4, containing 79 genes, is directed by its central hub, the bHLH transcription factor 036 - SlbHLH036 (*Solyc05g006650*). Among its co-expressed genes is the critical *Solyc03g122340.2* (Lipoxygenase D or LOX-D, highlighted in **Figure 3b**). This enzyme is vital in oxylipin production, influencing a variety of plant functions, from growth to defense. Specifically, LOX-D, part of the lipoxygenase pathway, oversees the creation of various oxylipins. The compounds from LOX-D are especially crucial in plant-pathogen dynamics and interplant signaling. The module accentuates LOX-D’s central role in producing oxylipins essential for plant defense and growth. Additionally, there’s a shift in gene activity from symptom emergence to the recovery stage, indicating its critical function in defending against PSTVd.

##### 3.1.1.3 Module 5: Cell Wall Dynamics and Divergent Strain Responses

Module 5, integrating 75 genes, is predominantly enriched in cell wall-associated processes. Two BHLH genes, *Solyc06g051550* and SlbHLH036, act as central connectors. The expression dynamics within this module exhibit strain-specific variations, especially between the two PSTVd strains. The M-strain showcases gene repression during heightened symptom manifestation, in contrast to the S-strain which depicts gene induction, underscoring differential strain responses. However, both strains evince a unison in gene induction during the recovery phase, alluding to shared recovery mechanisms post-infection.

#### 3.1.2 Modules that Exhibit Early Symptom Induction and Recovery Phase Repression in Response to the Severe Strain

These identified gene modules showcase a distinct pattern of gene expression in response to the severe strain of infection. During the early stages of infection, a synchronized induction of gene expression is observed, indicating the plant’s immediate response to the stressors induced by the severe strain of PSTVd. However, as the infection progresses and the plant enters the recovery phase, a notable shift occurs. Gene repression becomes prominent during this recovery stage, suggesting a dynamic regulatory mechanism at play. This intriguing phenomenon highlights the complex interplay between the plant’s defense mechanisms and its attempt to recover from the infection’s impact.

##### 3.1.2.1 Module 8: Unravelling Oxidative Stress Response and Lipid Biosynthesis Dynamics

Module 8, consisting of 49 genes, focuses on oxidative stress response and lipid formation. The hub gene, *Solyc03g031450*, networks with 15 distinct genes. Meanwhile, Palmitoyltransferase PFA4 (*Solyc10g086570*), linked with endoplasmic reticulum and Golgi apparatus formation, co-expresses with hub genes in this module. Throughout the infection by both PSTVd strains, a unified surge in gene expression occurs during pronounced symptom development, emphasizing the plant’s defensive measures against PSTVd stress. However, this amplification is counteracted by gene suppression during the recovery, hinting at a complex regulatory play in oxidative stress response and lipid biosynthesis in the context of PSTVd pathogenesis.

##### 3.1.2.2 Module 9: Unmasking Photosynthesis and Defense Response Dynamics

Module 9 comprises 43 genes linked to photosynthesis and defense. These genes exhibit varying activity between healthy and PSTVd-S infected plants. Specifically, there is an uptick in gene activity during early infection stages, which reverts during recovery. Specifically, the genes within this module display induction during the first and second stages of infection progression, aligning with symptom development. However, a notable departure from this pattern occurs during the recovery stage, characterized by gene repression. This significant shift in gene expression dynamics emphasizes the intricate interplay between photosynthesis and defense responses during PSTVd infections.In this module, five primary hub genes are identified in the root, all linked to photosynthesis. These include photosystem I reaction centre subunit V (*Solyc07g066150*); photosystem II 5 kDa protein, chloroplastic (*Solyc12g099650*); chlorophyll a-b binding protein 7, chloroplastic (*Solyc10g006230*) and two genes for Photosystem I reaction centre subunit XI (*Solyc06g082940* and *Solyc06g082950.2*). Their co-expression aligns with the module’s focus on photosynthesis and the photosystem. Additionally, *Solyc06g051550* is highlighted as a key regulator in this module, similar to its role in other modules.

### 3.2. Exploring Leaf Tissue Gene Co-Expression Modules

Shifting our focus to leaf tissue, our exploration highlights gene co-expression patterns that elucidate the plant’s molecular reactions in the presence of PSTVd. Our research identifies seven distinct gene co-expression modules (M1 to M7) within the leaf tissue. These modules, ranging in size from 24 to 331 genes, embody the complex interplay of genes and regulators reacting to the pathogen. Specifically, modules M5, M1, M3, M6, and M2 emerge as particularly noteworthy due to their pronounced regulatory shifts across conditions, underlining their essential roles in the plant’s defensive responses to PSTVd infections.

#### 3.2.1 Shared Regulatory Behaviour: A Consistent Pattern

Notably, four of these intriguing modules—M5, M1, M3, and M6—exhibit a remarkably similar pattern of behaviour. In control plants, a consistent trend of induction is observed across almost all stages of study. This underscores the essential role of these modules in normal cellular processes and homeostasis. However, an intriguing contrast emerges when examining PSTVd-S infected plants. These modules, which normally show induction in control plants, exhibit a notable trend of repression across the majority of studied stages. This observation hints at a nuanced interplay between PSTVd-S infection and the plant’s regulatory machinery, leading to distinct modulation of these modules, **Figure 6**.

**Figure 4.**
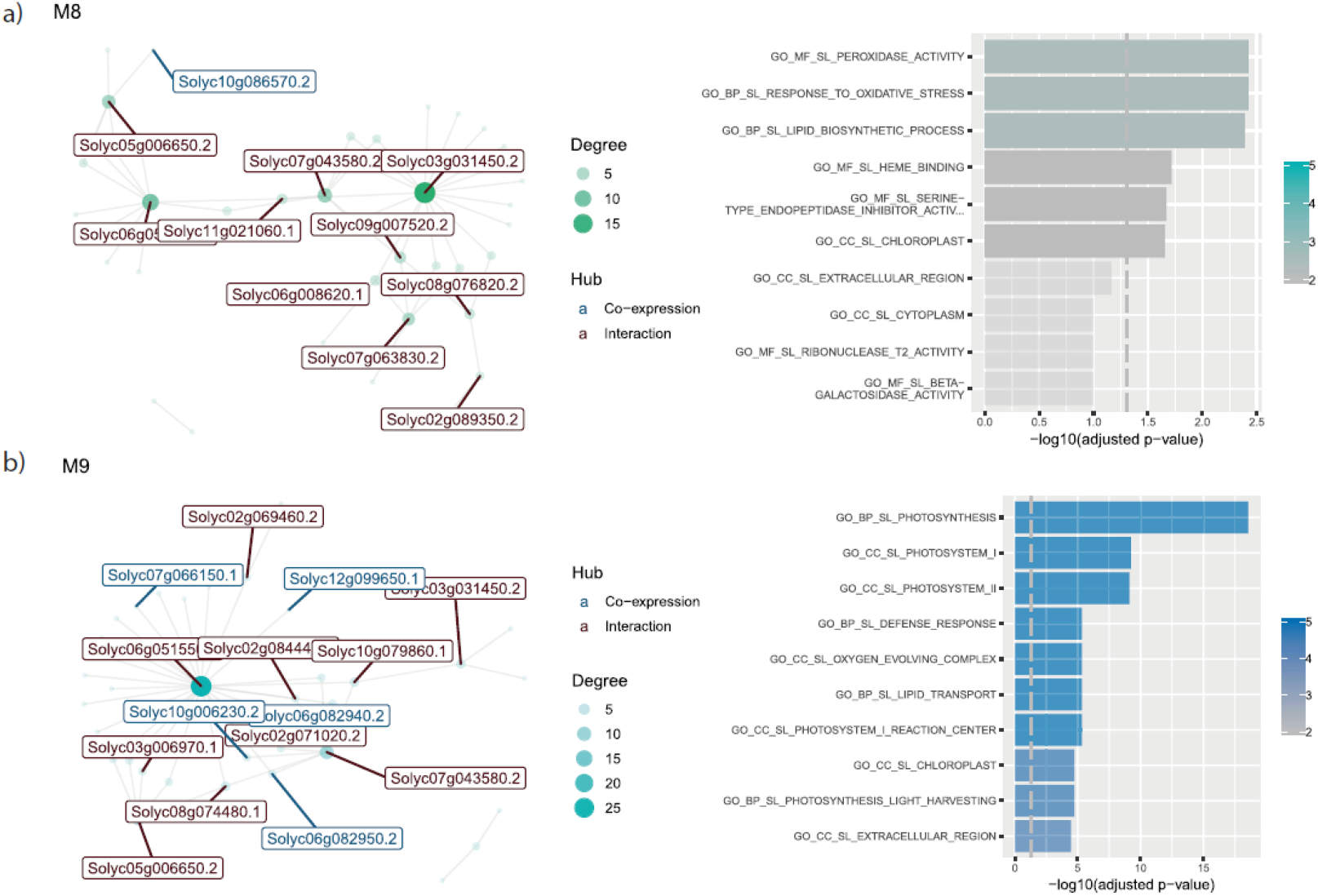
Co-expression interaction network and functional enrichment of modules at early symptoms and repressed at recovery in root-tissue. Each node on the network represents a gene, size is proportional to the amount of interactions (degree). Hub genes are listed with their names, in red if they only interact, and in blue if they are co-expressed with their targets. Next to each network, biological functions of their genes are shown. Length of the bar represents significance, and color gradient number of genes implicated in each Gene Ontology Term. a) Interaction network and biological functions for M8; b) Interaction network and biological functions for M9.

**Figure 5.**
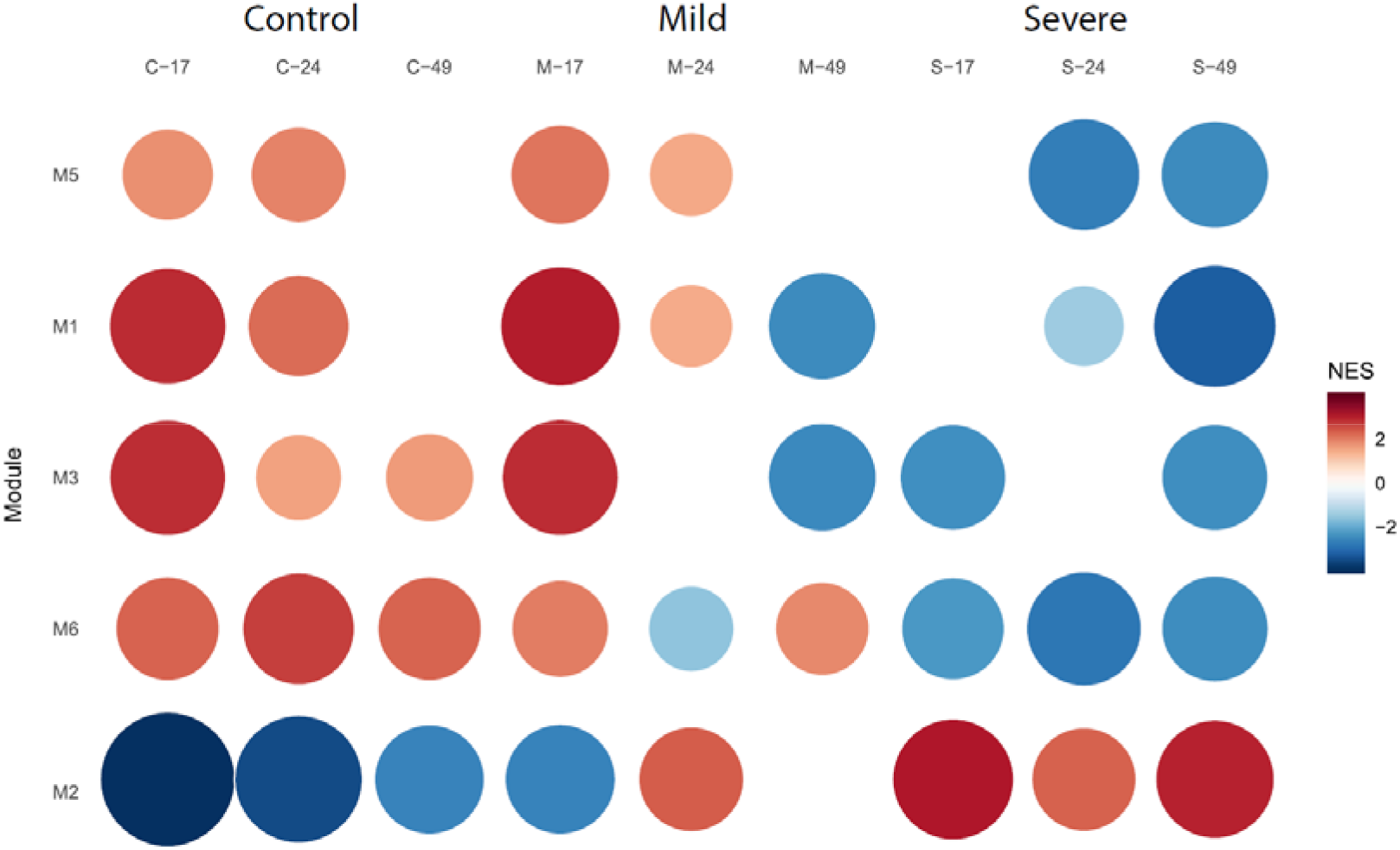
Dynamics of Co-Expression Modules in Leaf Tissue. GSEA for the significant co-expression modules (M5, M1, M3, M6, and M2) within leaf tissue. It visually represents the module activity under control, PSTVd M, and PSTVd S23 strains during the time course conditions. A visual representation captures the regulatory dynamics of the five key co-expression modules (M5, M1, M3, M6, and M2) within leaf tissue. The figure illustrates the contrasting behaviour of these modules in control plants and those infected with PSTVd-S. The graph underscores the intricate interplay of induction and repression across different stages, unravelling the complex regulatory mechanisms that govern the plant’s response.

**Figure 6.**
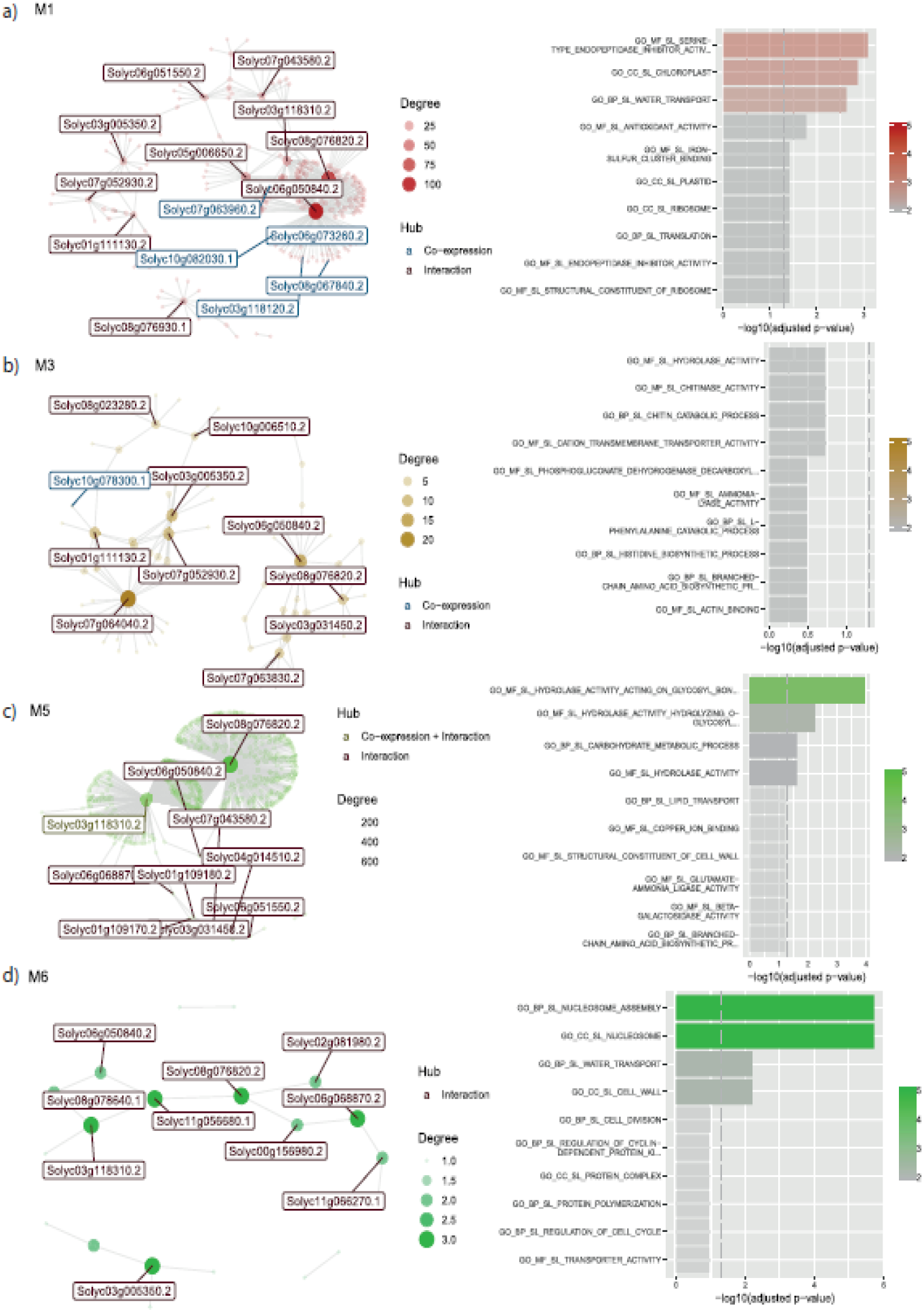
Co-expression interaction network and functional enrichment of modules repressed at recovery with PSTVd-S23 in leaf. Each node on the network represents a gene, size is proportional to the amount of interactions (degree). Hub genes are listed with their names, in red if they only interact, and in blue if they are co-expressed with their targets. Next to each network, biological functions of their genes are shown. Length of the bar represents significance, and color gradient number of genes implicated in each Gene Ontology Term. a) Interaction network and biological functions for M1; b) Interaction network and biological functions for M3; c) Interaction network and biological functions for M5; d) Interaction network and biological functions for M6.

##### 3.2.1.1 Module 1: Orchestrating Chloroplast Biogenesis and Beyond

Module 1, comprising 331 genes, captures our attention with its intricacy and significance. This intricate network has five co-expressed genes—*Solyc07g063960.2* (50S ribosomal protein L24), *Solyc06g073260.2* (NAD-dependent epimerase/dehydratase), *Solyc10g082030.1* (Peroxiredoxin), *Solyc08g067840.2* (PsbP domain-containing protein 5, chloroplastic), and *Solyc03g118120.2* (Transferase transferring glycosyl groups). These genes exert influence over a range of interactions, spanning from 25 to 100, highlighting their pivotal roles within the module. Additionally, SlbHLH135 (*Solyc06g050840*) is a hub gene having more than a hundred interactions. The genes within Module 1 encompass a diverse array of functions, shedding light on the multifaceted roles they play within leaf tissue. In chloroplast biogenesis, these genes contribute to vital processes, ensuring the plant’s capacity to harness light energy for growth and sustenance. Additionally, their involvement in water transport and protein translation unveils the interwoven processes that underlie the leaf’s dynamic response to PSTVd infection.

##### 3.2.1.2 M3: Exploring the Landscape of Metabolic Regulation

Within Module 3, comprising 134 genes, parallels can be drawn with Module 1’s behavioural dynamics. Central to this module is the co-expressed hub-gene *Solyc10g078300.1*, associated with single-stranded nucleic acid binding R3H protein, emphasizing its role in metabolic regulation. Moreover, the Photosystem I reaction centre subunit V (*Solyc07g064040*) emerges as a pivotal regulator, interacting extensively with over 20 distinct genes. The influence of this gene assembly extends to various biomolecules such as chitinase, phenylalanine, and histidine, underscoring the multifaceted metabolic adaptations prompted by pathogen invasion.

##### 3.2.1.3 M6: Navigating Cell Division and Beyond

Module 6, with 28 co-expressed genes, navigates the critical landscape of cell division. Its genes, entwined in processes such as nucleosome assembly, nucleosome as a cellular component, and the very essence of cell division, unveil a tightly orchestrated performance within the leaf tissue. The behaviour of this module underscores the leaf’s determination to adapt and thrive in the face of adversity.

#### 3.2.2 Module 2: A Dynamic Contrast in Defense

In contrast to other modules, Module 2, with 149 genes, exhibits a unique response to PSTVd infection. Furthermore, a hub gene with more than 600 interactions is shown for this module: SlbHLH135 (*Solyc06g050840*). While other modules are induced, genes in Module 2 are repressed in control samples, suggesting a readiness for defense. However, during PSTVd-S infection, a strong gene activation occurs across multiple stages. This module primarily emphasizes the leaf’s defense mechanism against biotic threats. The gene ontology terms for Module 2 reveal the leaf’s commitment to bolstering its defenses against pathogens, highlighting the e dynamics of plant-pathogen interactions and the leaf’s adaptive capacity.

### 3.3. Bipartite Networks: Unveiling Specific Regulatory Patterns

To enhance our comprehension of how gene regulation occurs, we constructed separate bipartite tissue-specific networks characterizing the regulation of target genes by all the bHLH TFs in each network. These bipartite networks consist of two layers: one layer of regulators, encompassing all the genes corresponding to bHLH TFs, and a second layer of target genes, encompassing all the genes in the network regulated by a bHLH TF. For instance, the bipartite network of the leaf comprises 30 bHLH genes, whereas the root network contains 28 bHLH genes. Both networks have regulatory guidance from a common set of 28 bHLH genes. In other words, the leaf network possesses the same 28 bHLH genes found in the root network but also includes two additional bHLH genes, specifically: *Solyc03g097820.1* SlbHLH022 involved in physiological, developmental and metabolic processes; and the drought responsive protein encoded by *Solyc06g072520.1* GBOF-1.

#### 3.3.1. Insights from the Root Bipartite Network

For root samples, the bipartite network comprises 28 bHLHs and 1004 target genes. This network reveals 563 genes regulated by one bHLH, 316 targets regulated by two bHLHs, 93 targets regulated by three bHLHs, and 32 targets regulated by four bHLHs—none of the genes are regulated by more than four bHLHs. Functional enrichment analysis sheds light on the roles of these regulated genes. Genes with a single regulator are implicated in *Chloroplast, Photosystems I and II*, while genes regulated by two bHLHs play a part in *KEGG 00592 Alpha-linoleic acid metabolism*, **Figure 8**. For instance, the bHLH-Jasmonic Acid 3 *Solyc08g076930.1* (IA3) is a key regulator within this network, controlling the plant’s response to pathogens through the accumulation of jasmonic acid, a primary defense mechanism activated by plants upon pathogen infection.

**Figure 7.**
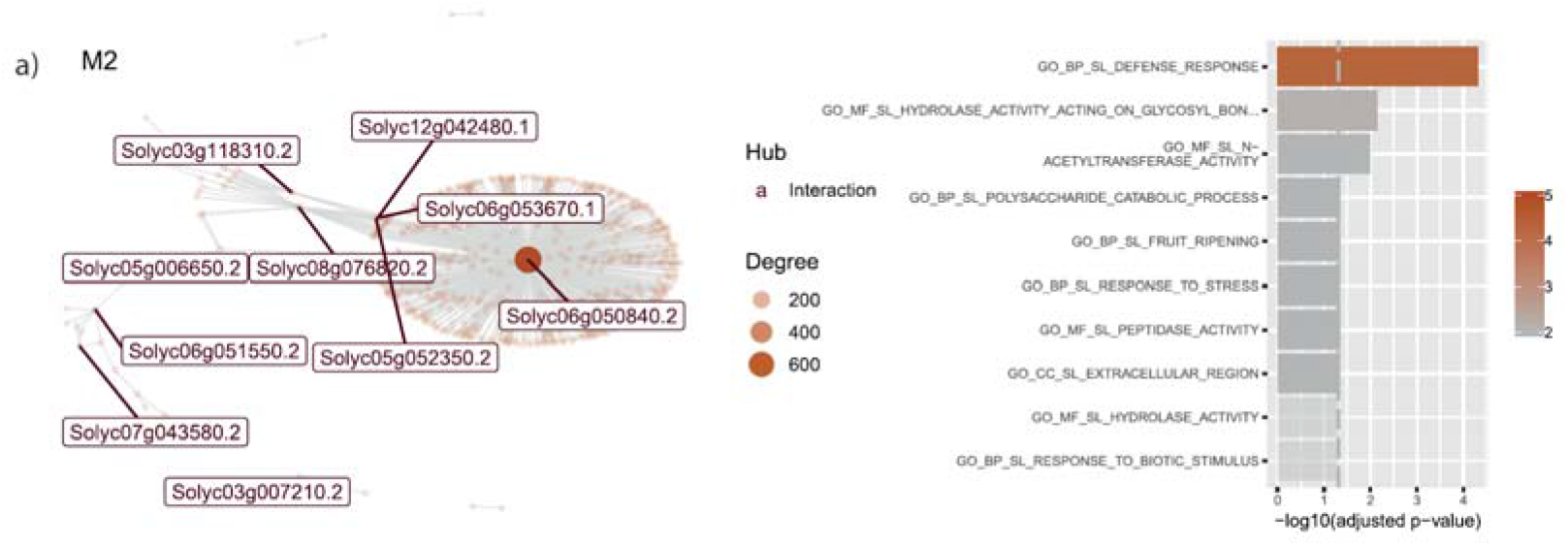
Co-expression interaction network and functional enrichment of module 2 in leaf. Each node on the network represents a gene, size is proportional to the amount of interactions (degree). Hub genes are listed with their names, in red if they only interact, and in blue if they are co-expressed with their targets. Next to each network, biological functions of their genes are shown. Length of the bar represents significance, and color gradient number of genes implicated in each Gene Ontology Term. a) Interaction network and biological functions for M2.

**Figure 8.**
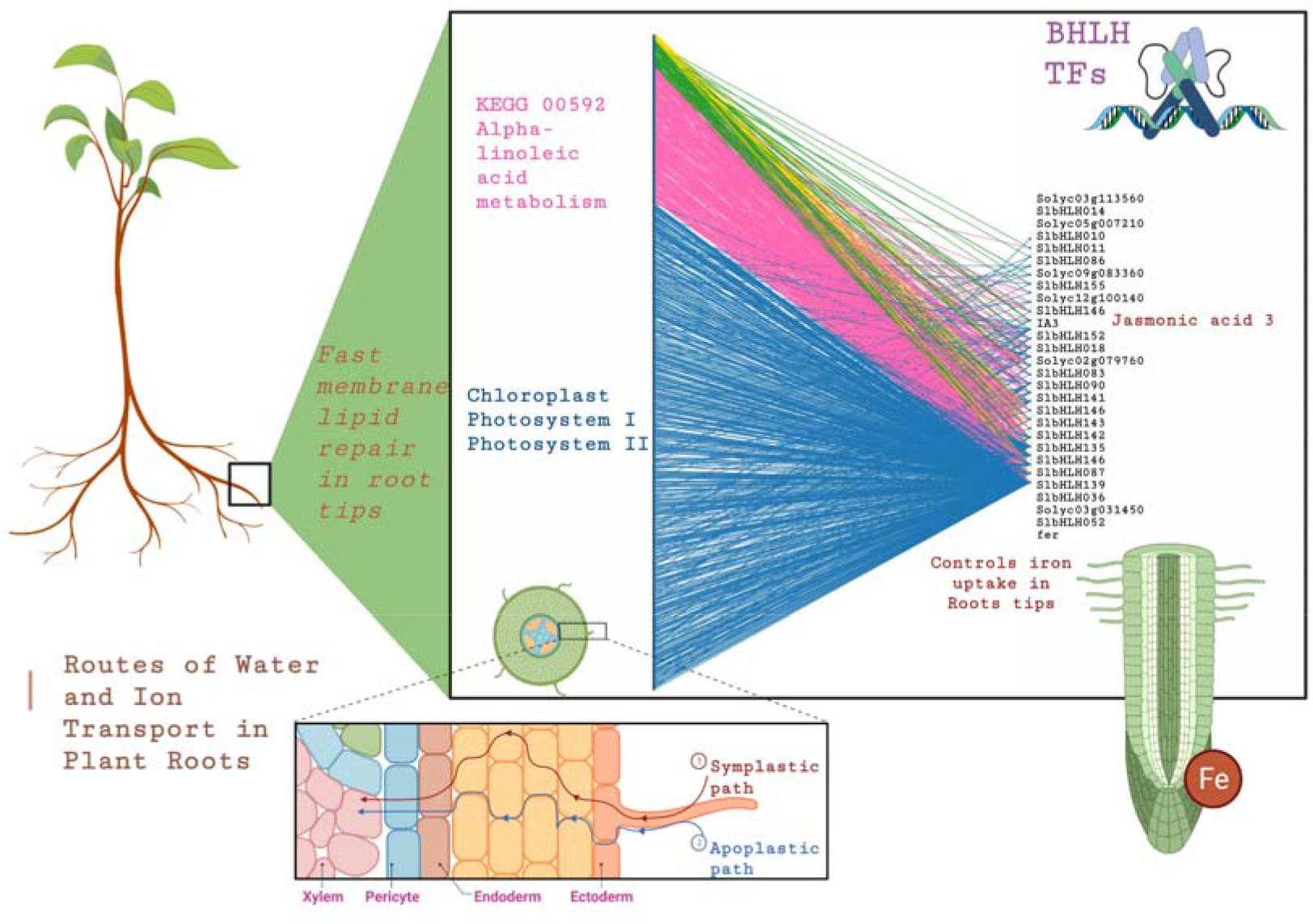
Deciphering Root-Specific Regulatory Patterns. **a)** bipartite network analysis unveils connections between bHLH (right) TFs and target genes (left). Connections-colours represent the number of bHLH regulation of each target gene as follows: blue for one, pink for two, green for three, and yellow for four. While functional enrichment analysis highlights specific pathways, including chloroplast, Photosystem I and Photosystem II, and KEGG 00592 Alpha-linoleic acid metabolism.

#### 3.3.2 Insights from the Leaf Bipartite Network

The leaf bipartite networks identify interactions between 30 bHLHs and 1970 target genes. These genes are categorized based on the number of TFs regulating them: 715 targets regulated by a unique bHLH, 874 targets regulated by two bHLHs, 336 targets regulated by three bHLHs, 40 targets regulated by four bHLHs, and 5 targets regulated by more than four bHLHs. This regulatory behaviour of more than four regulators is only present in the leaf tissue-specific network. Similar to its presence in the root, *Solyc08g076930.1* IA3 is also a regulator in the leaf network, underscoring its significance as a key player in plant responses to pathogen infections. A functional enrichment analysis for these five groups was conducted. We found that the *KEGG:00073 Cutin, suberine, and wax biosynthesis* is regulated by unique TFs, while the aspartic-type endopeptidase activity has two regulators, and ornithine metabolic processes has genes with three regulators, **Figure 9**.

**Figure 9.**
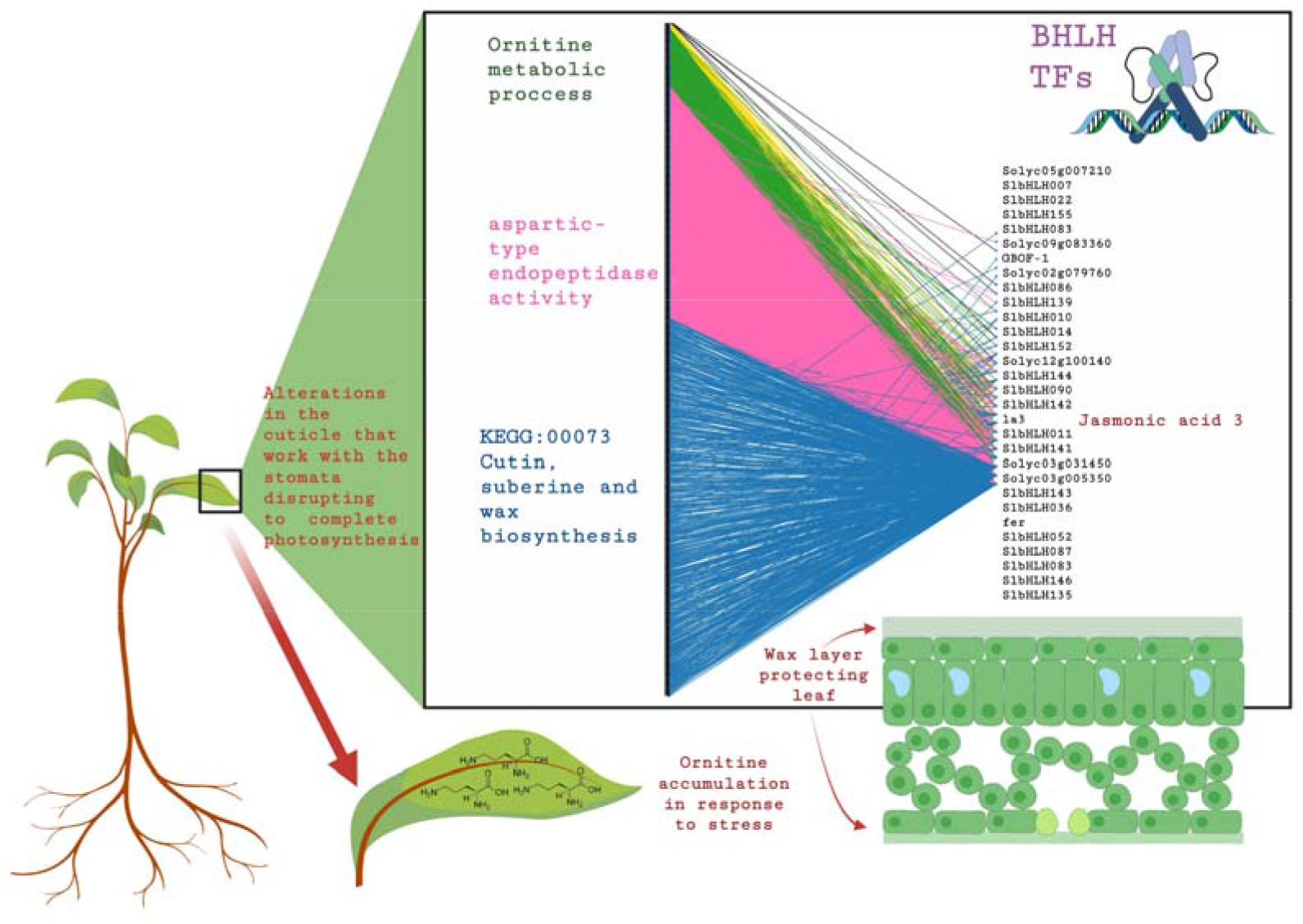
Deciphering Leaf-Specific Regulatory Patterns. a) bipartite network analysis unveils connections between bHLH (right) TFs and target genes (left). Connection color represents the number of bHLH regulation of each target gene as follows: blue for one, pink for two, green for three, and yellow for four. While functional enrichment analysis highlights specific pathways, including cutin, suberine, and wax biosynthesis, endopeptidase activity, and ornithine metabolic processes, the leaf-specific network showcases the exclusive presence of genes regulated by more than four TFs.

### 3.4. Mapping the Regulatory Landscape: A Bridge Between Modules

Within the complex landscape of gene interactions highlighted in our study, the bifan motif stands out as a prominent regulatory beacon. This motif, a distinctive architecture of regulatory interactions, plays a central role in deciphering the sophisticated governance of gene regulation within our co-expression modules. The bifan motif is emblematic of a specific regulatory paradigm, distinguished by its two primary *“source”* nodes, represented by bHLH TFs, and two secondary *“target”* nodes, which are the genes under regulation, **Figure 10**.

**Figure 10.**
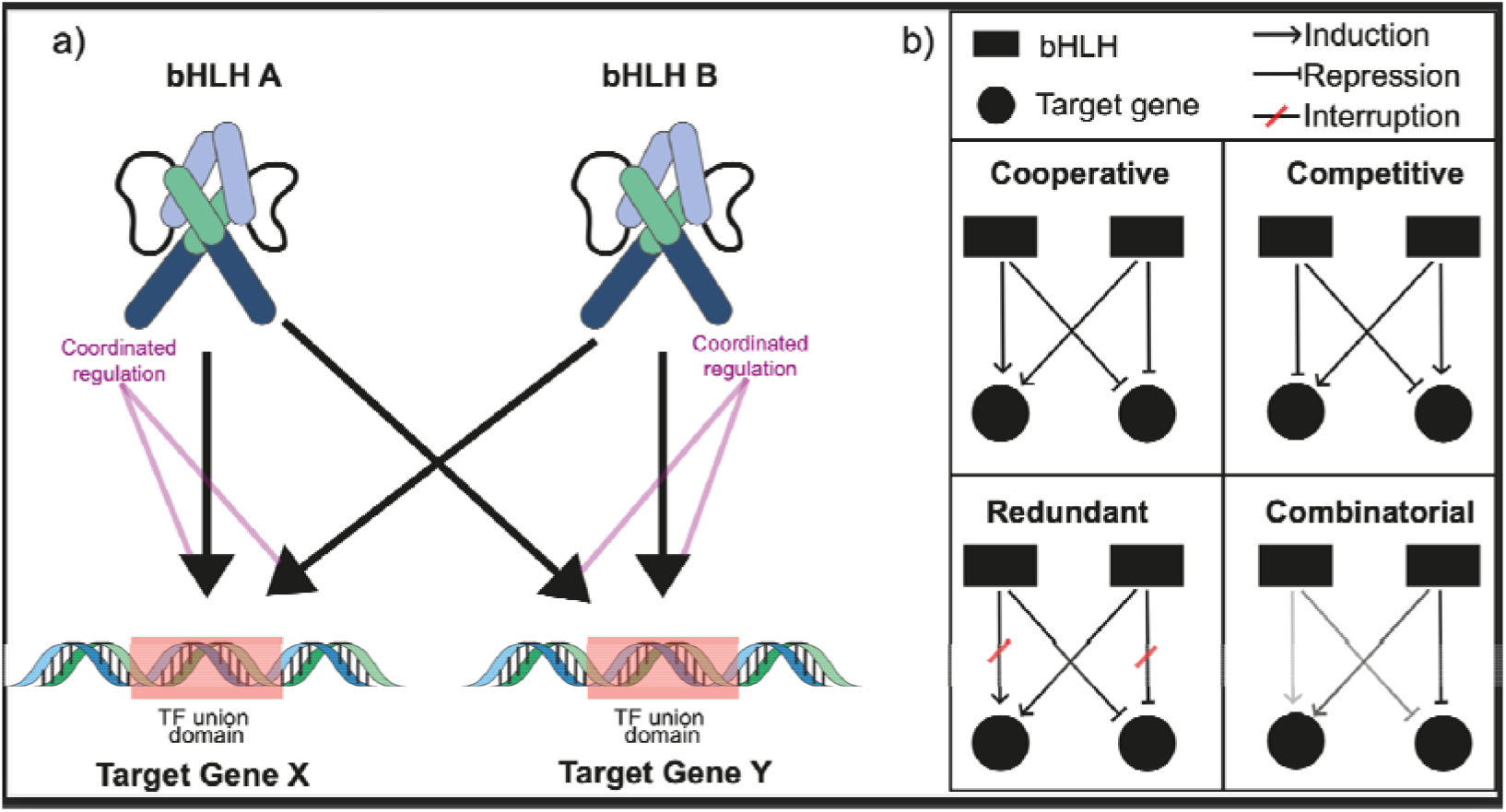
Bifan Motif and its functionality in gene regulation. a) Graphic illustration of the Bifan motif: two different bHLH regulating the expression of two different target genes; arrows depict regulation; b) Examples of Bifan motif activity in gene regulation. Legend for the diagram is shown at the top. Four different regulation types are shown: Cooperative, Competitive, Redundant, and Combinatorial. Color gradient indicates level of regulation: the darker, the stronger.

A defining feature of this configuration is the independent regulation by each source node of both target nodes. The structure of the bifan motif enables a coordinated regulatory response, wherein alterations in the activity or levels of nodes A and B correspondingly modulate the expression or activity of nodes X and Y, as visualized in **Figure 10a**. Beyond its unique dual-regulatory characteristic, the bifan motif facilitates an intricate synergy of transcriptional control within our identified gene modules. Crucially, this motif serves as a linchpin, bridging separate modules and revealing convergent and divergent regulatory patterns that intricately weave through the vast tapestry of gene interactions. The bifan motif underscores several pivotal attributes such as *Flexibility:* the dual regulatory nodes confer a degree of adaptability, enabling the system to recalibrate in response to nuanced environmental changes, *Robustness:* built-in redundancy ensures resilience against disruption, safeguarding the integrity of regulatory processes and *Signal Integration:* adept at amalgamating signals from disparate pathways, enabling the cellular machinery to craft a unified and coherent response, **Figure 10b**. The prevalence of the bifan motif within our co-expression networks deciphers the regulatory intricacies governing the plant’s response to PSTVd infection. This motif’s role as a nexus of cross-talk underscores its significance in shaping the complex interplay of gene expressions, shedding light on the molecular dialogues that govern the defense and adaptation strategies of the host plant.

## Discussion

Given the complexity of plant-pathogen interactions, dissecting the regulatory framework is of paramount importance. Our study delves deep into gene co-expression modules, shedding light on the molecular intricacies of the host during both mild and severe PSTVd infections across root and leaf tissues.

Root tissue analysis reveals bHLH guided key regulatory processes such as – heme-binding (M2), ubiquitin defense (M4), and cell-wall processes (M5) – pivotal during symptom development due to the severe strain. These biological events synchronize with robust gene expression, emphasizing the plant’s heightened defense. This synchronization is an evolutionary response to aggressive pathogens. Notably, hub genes emerge as the linchpins, much like regulatory keystones, ensuring the efficiency of defense strategies. Their dense interconnectivity suggests roles as signal integrators and response amplifiers. Specifically, *Solyc06g051550*-heme binding, alongside *Solyc06g051550* and SlbHLH036, prove critical during the severe symptom manifestation.

For instance, Module 2 emphasizes *Solyc06g051550*-heme binding role with the hub gene UDP-glycosyltransferase 73C3 in glycosylation processes. UDP-glycosyltransferases, essential for sugar attachment to substrates, oversee various cellular operations. This gene’s vast co-expression connections affirm its central role in glycosylation functions, which is fundamental for the plant’s defense during infection. Additionally, *Solyc06g051550* and SlbHLH036’s influence on ubiquitin defense related to Lipoxygenase D. This enzyme plays a key role in producing oxylipins, precursors to jasmonic acid, which instigates defense mechanisms against pathogens.

Early symptom induction and the subsequent recovery phase in root tissue accentuate the relevance of bHLH TFs like *Solyc03g031450* and *Solyc10g086570*. Additionally, the synergy between these genes and those related to photosynthesis, namely *Solyc07g066150, Solyc12g099650, Solyc10g006230, Solyc06g082940,* and *Solyc06g082950.2*, suggests a host protective strategy against viroid effects, potentially preserving energy production. For leaf responses, SlbHLH135 (*Solyc06g050840*) stands out due to its critical function in chloroplast formation and defense. Additionally, *Solyc10g078300.1* has been identified as integral for metabolic stability under stress. Meanwhile, Module 5 is centred around the bTF SlbHLH083 hub gene, intricately interacting with numerous other genes. These interactions orchestrate vital processes such as hydrolase activity and carbohydrate metabolic processes. Given its extensive reach within the network, this BbLH transcription factor exerts control over a variety of cellular functions, thereby contributing to the plant’s adaptability and response to PSTVd infection through coordinated metabolic and energy-related processes. Moreover, module 1 tissue is guided by a group of hub genes, including 50S ribosomal protein L24, NAD-dependent epimerase/dehydratase, peroxiredoxin, PsbP domain-containing protein 5 (chloroplastic), and transferase transferring glycosyl groups. These hub genes collectively oversee a wide spectrum of functions encompassing chloroplast biogenesis, translation, and transferase activity. Their orchestrated regulatory roles influence processes vital for the plant’s survival, including energy production and structural integrity.

To summarize, the comparative analysis of gene co-expression modules and their regulatory patterns in leaf and root tissues unveils tissue-specific responses to severe PSTVd strain infection. While certain modules demonstrate shared behaviours, such as robust induction during symptom development, others display distinctive regulatory patterns, shedding light on the unique roles played by each tissue in the plant’s defense and adaptation strategies. Further analysis of the bipartite networks specific to each tissue reveals varied regulatory patterns based on the presence of bHLH regulators. Root tissue focuses on processes like photosynthesis, while leaf tissue showcases unique patterns related to structural integrity and defense. Moreover, 28 bHLHs genes recur in both networks, hinting at a universal defense strategy. For instance, The bHLH-Jasmonic Acid 3 *Solyc08g076930.1*(IA3) regulator plays a pivotal role in plant defense against pathogens, primarily through its connection with the jasmonic acid (JA) pathway. JA is a crucial defense molecule that activates upon pathogen detection, signaling other defense mechanisms and producing deterrents against herbivores and pathogens. The IA3 regulator enhances the efficiency of this response by potentially modulating the intensity or longevity of JA’s action, ensuring optimal defense against varying threats. The bipartite analysis amplifies the bHLH regulatory theme, highlighting specific defense mechanisms in the leaf network regulated by *Solyc03g097820.1* SlbHLH022 and drought-responsive *Solyc06g072520.1.GBOF-1*.

In accordance with our previous research, the role of those bHLH-TFs (*Solyc08g076930.1* IA3,*Solyc03g097820.1* SlbHLH022 and *Solyc06g072520.1.GBOF-1*.) becomes even more compelling [38]. Having identified them as a master transcriptional regulators during mild and severe strain infections respectively, their reappearance in the present analysis acts as a reaffirmation of their importance in the defense of the plant. Such consistency in findings across different studies points to the reliability and robustness of those genes as key players in plant defense in the PSTVd-tomato pathosystem.

The recurrent presence of the bifan motif underscores its crucial role as a central player in coordinating these responses, offering a comprehensive comprehension of the intricate molecular dialogues that unfold in response to pathogenic challenges. In conclusion, bHLH-TFs within co-expression modules play a central role in managing complex molecular interactions during severe PSTVd infections in both root and leaf tissues. These TFs guide the regulation of indispensable biological roles spanning from glycosylation and oxylipin biosynthesis to chloroplast biogenesis and translation. Their influence extends beyond individual processes, promoting communication and coordinated responses. The identification and characterization of the hub genes enhance our understanding of the regulatory landscape, offering insights into the plant’s strategies for defense and adaptation against the challenges posed in the contrasting PSTVd-tomato pathosystem.

bHLH TFs are key regulators in pospiviroid infections, transcending their roles in leaves and roots. In floral tissues, they exert a marked influence on petal morphology. A case in point is the tomato gene, SlBIGPETAL1 (*Solyc05006650.2.1*), which governs petal architecture and mirrors the role of its Arabidopsis counterpart [At1g59640.2], [47]. This reflects the evolutionary preservation of bHLH factors, emphasizing their versatility in modulating diverse plant anatomies across various tissues. Moreover, their role extends far beyond the tomato-PSTVd pathosystem, as they serve as pivotal regulatory elements in various plant-pathogen interactions. In hops, these factors influence the production of key metabolites, like flavonoids and bitter acids, critical for brewing. Specifically, the HlMYB7 bHLH factor suppresses genes related to these metabolites, impacting plant growth, flowering, and viroid resistance [48]. In sweet cherry, post HSVd infection, there’s a noticeable change in bHLH genes, notably the upregulation of bHLH79, affecting fruit health and productivity [49]. These insights highlight the bHLH factors’ central role in plant defense and metabolism, suggesting their potential in enhancing plant resilience and metabolic efficiency.

### Conclusions

Viroid infections prompt complex genetic reprogramming in host plants, with bHLH transcription factors orchestrating defense gene activation while limiting viroid replication. bHLH TFs exhibit tissue-specific expressions, adapting defense strategies accordingly and interacting with other regulatory proteins to orchestrate context-specific responses. Gene modules under bHLH TF control encompass a range of functions from defense to stress responses and metabolic adjustments. Hormonal pathways like jasmonic acid intersect with bHLH TFs, highlighting the intricate nature of the plant’s defense mechanisms. Unresolved questions persist, urging deeper exploration into the mechanisms through which bHLH transcription factors guide the host’s response to viroid infection. This pursuit offers essential insights for strategies to enhance plant resistance.

## Acknowledgements

This research was funded by internal USDA-ARS project number 8042-22000-318-00D. K.A.-P. (CVU:227919), O.Z.-M. (CVU:1147042) and M.A.J.-L. (CVU:1035685) received financial support from CONAHCyT. K.A.-P. had a fellowship from the Fulbright García-Robles foundation.

